## Supplementary figures and tables for "Cyclic di-AMP traps proton-coupled K^+^ transporters of the KUP family in an inward-occluded conformation"

### Supplementary information

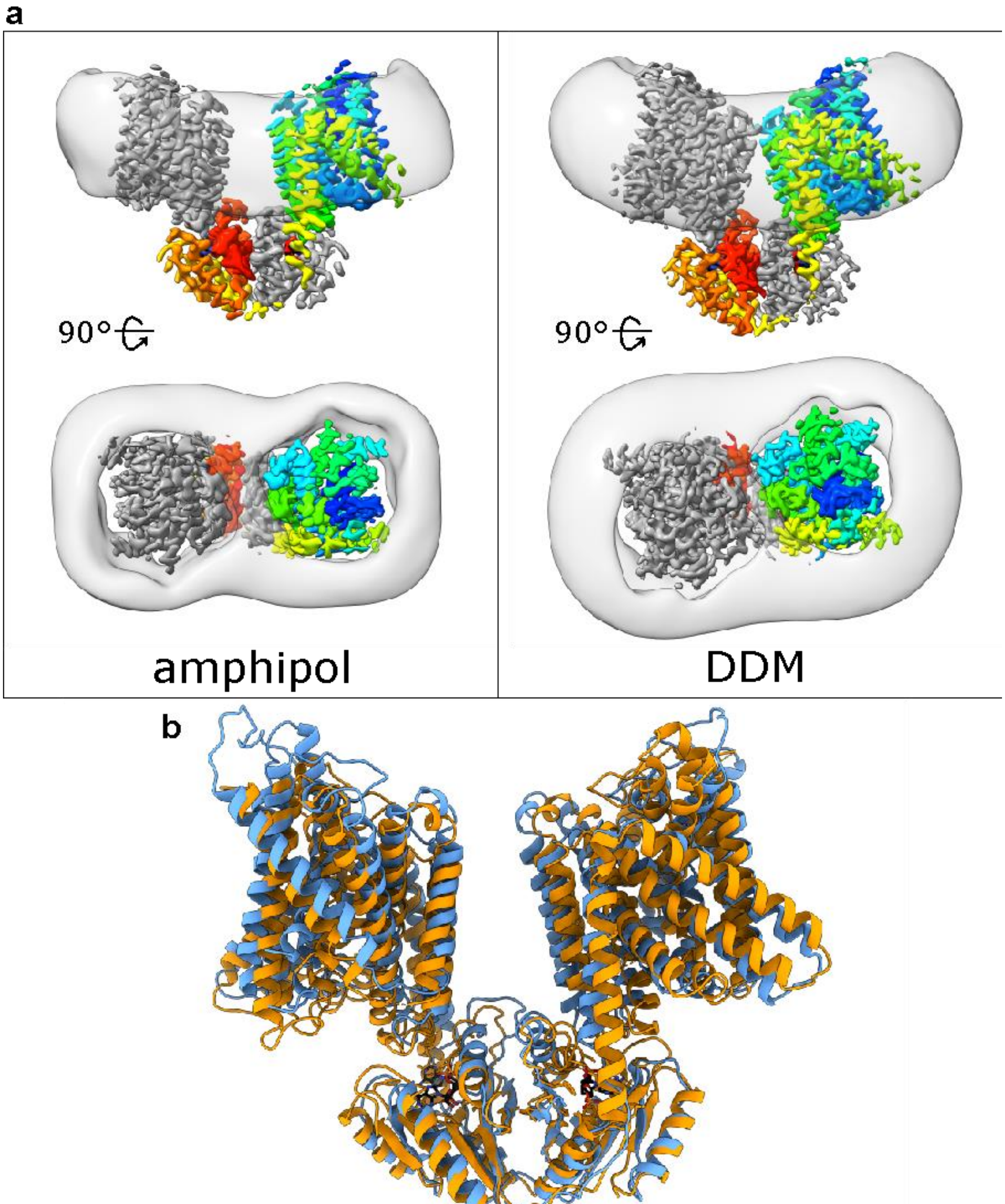


**Supplementary Figure 1: a**) 3.8 Å and 3.3 Å maps of KimA with gaussian filtered ( ʃ = 3) amphipol and DDM belt, respectively. The relaxed, upright dimer architecture of KimA is preserved in the amphipol belt that forms around each TMD (left). In DDM, the TMDs tilt inwards as previously seen in SMALPs. Density is coloured by monomer. One monomer is coloured in a rainbow scheme from N to C terminus. **b**) View of amphipol-reconstituted KimA (orange) aligned on cytoplasmic residues 462 to 606 with the upright dimer as previously predicted using MD simulations[25] (blue). The MD structure was constructed by clustering ca. 6.2 µs of atomistic MD data using gmx cluster with an RMSD cutoff of 0.3 nm. Shown is the state which best represents the cluster for the upright dimer, which accounts for approximately 33% of the total data.

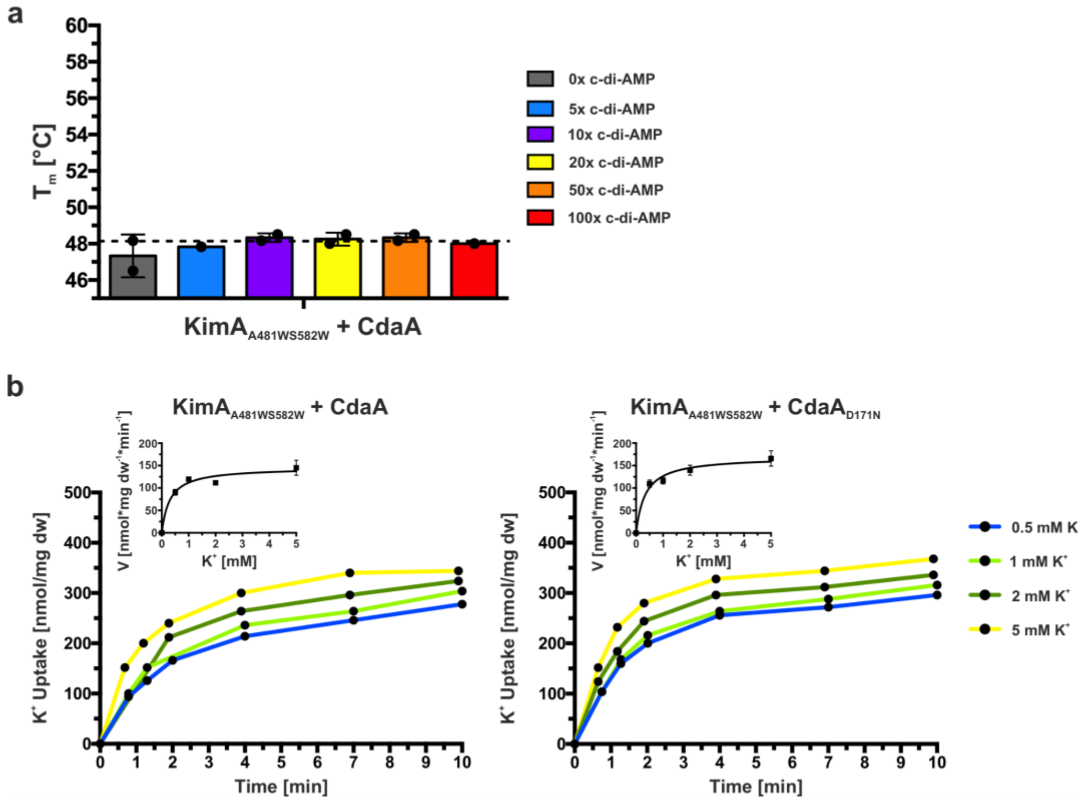


**Supplementary Figure 2: Abolished inhibition by c-di-AMP due to steric blockage of binding pocket in KimA$_{A481WS582W}$ variant.** **a)** Melting temperature of KimA$_{A481WS582W}$ purified from cells with CdaA present (KimA$_{A481WS582W}$ + CdaA) and incubated with an increasing c-di-AMP concentration given in x-fold molar excess over KimA. Determined with Differential Scanning Fluorometry (DSF). Dashed line indicates $T_m$ of KimA WT w/o c-di-AMP addition. Data points represent the average and error bars the standard deviation of measurements from a biological duplicate. **b)** Potassium uptake of *E. coli* LB2003 cells expressing *kimA$_{A481WS582W}$* and *cdaA*/*cdaA$_{D171N}$* at increasing external potassium concentrations. One representative experiment shown (n= 4/4). Michaelis-Menten diagram included in graph.

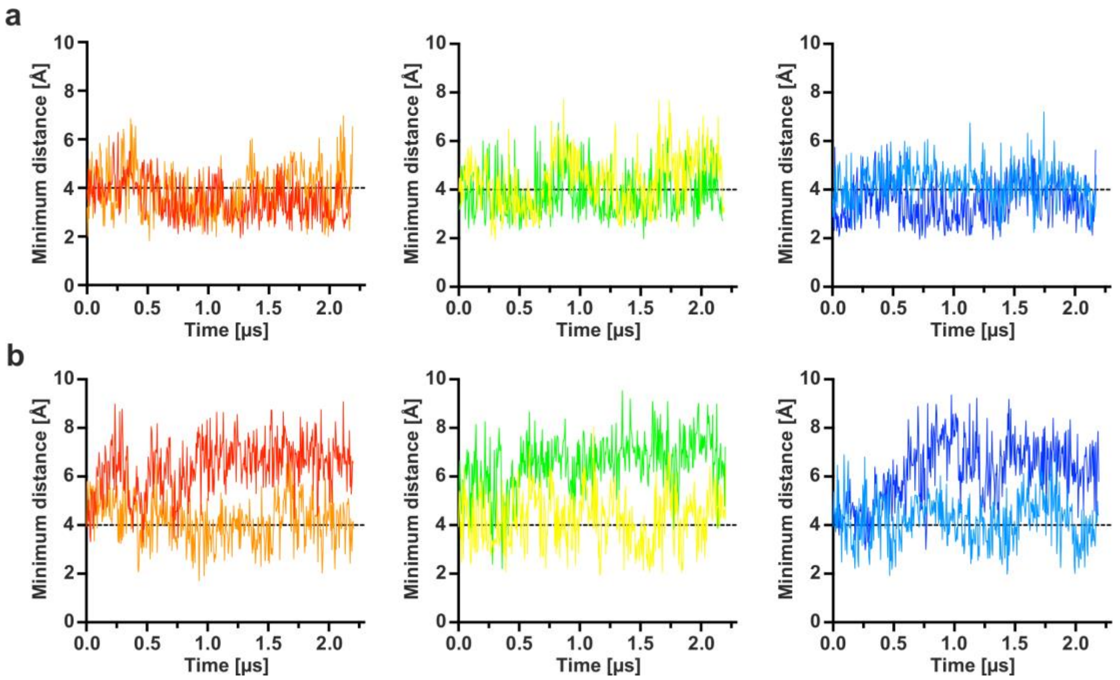


**Supplementary Figure 3: Minimum distance plots between Tyr118 and Asn237 with ligand (a) and without ligand (b).** Each graph is for one simulation, with the two traces representing the two KimA monomers. A dotted line at 4 Å denotes the approximate cutoff for whether the residues are in contact.

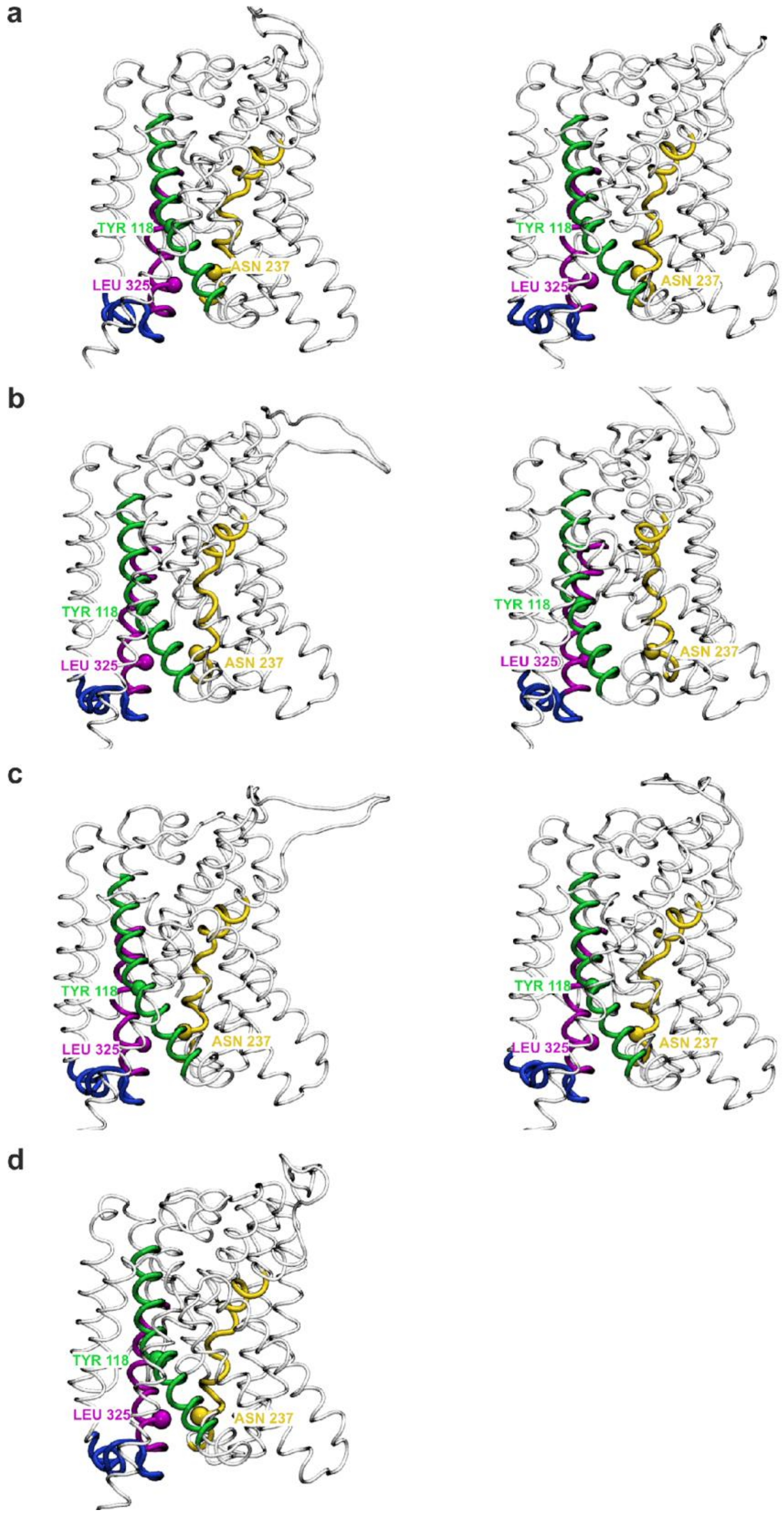


**Supplementary Figure 4: Eigenvector 1 and 2 extreme views**. As Figure 5c, eigenvector 1 extreme views but for the ligand-bound state (a), as well as eigenvector 2 for both states (b: no-ligand, c: ligand) and the relative view from the input structure (d). As in eigenvector 1, in eigenvector 2 there is movement of TM6 away from TM3/8 in the non-liganded state. TMH6 (yellow), TMH3 (green) and TMH8 (pink), loop connecting TMH8 and 9 (blue).

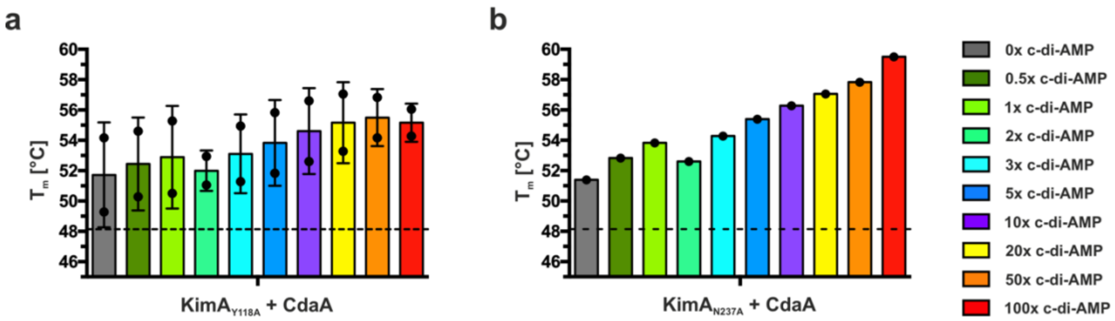


**Supplementary Figure 5: Melting temperature of KimA$_{Y118A}$ + CdaA (a) and KimA$_{N237A}$ + CdaA (b).** Melting temperatures of KimA variants purified from cells with CdaA and incubated with an increasing c-di-AMP concentration given in x-fold molar excess over KimA. Determined with Differential Scanning Fluorometry (DSF). Dashed line indicates T$_m$ of KimA WT w/o c di AMP addition.

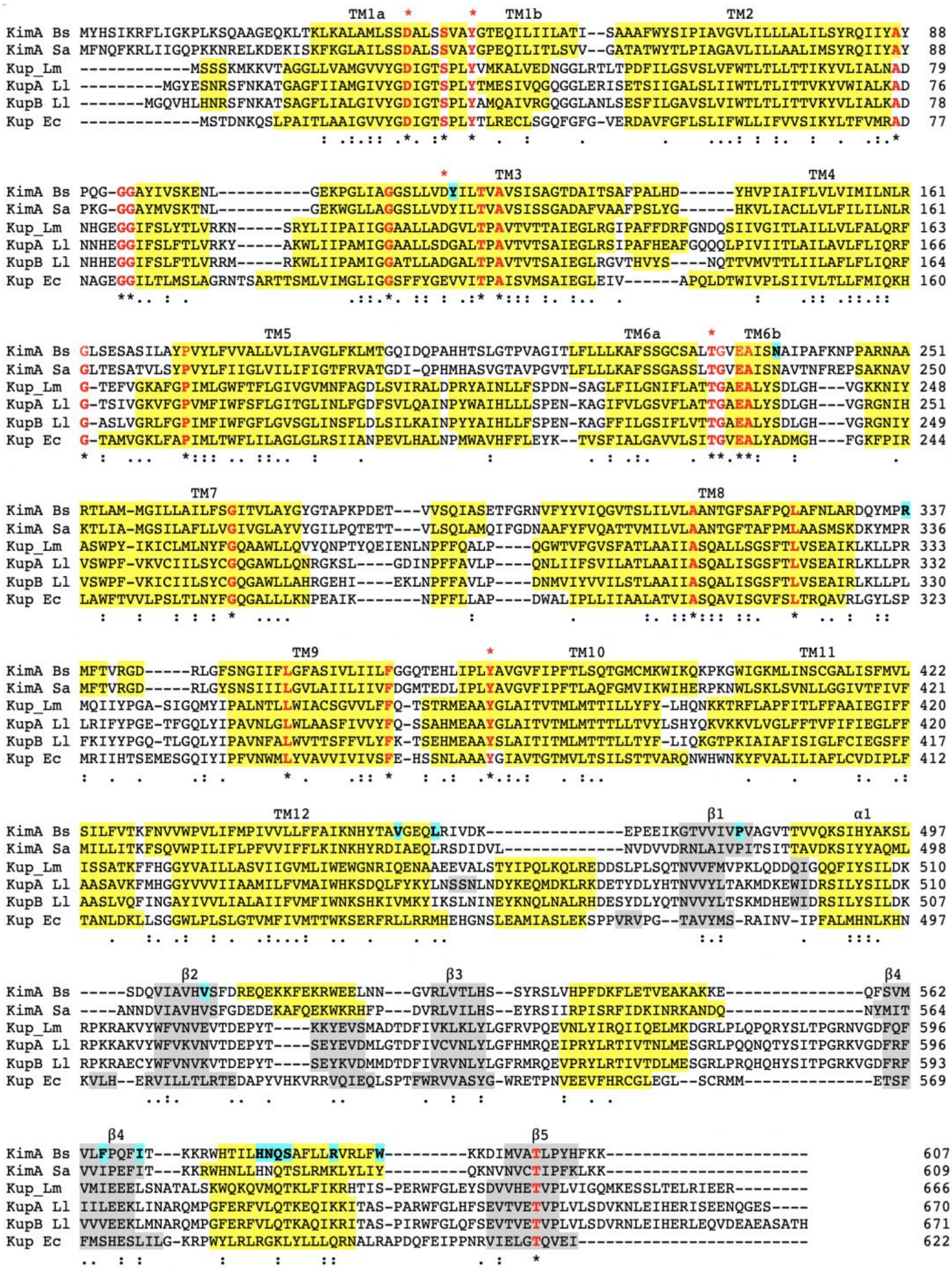


**Supplementary Figure 6: Structure-based alignment of KUPs.** AlphaFold predictions of KimA from *S. aureus* (KimA Sa), Kup from *L. monocytogenes* (Kup Lm), KupA and KupB from *L. lactis* IL1403 (KupA Ll and KupB Ll), and Kup from *E. coli* (Kup Ec) were aligned to the *B. subtilis* KimA structure (KimA Bs). Conserved residues are in bold red font, α-helices are highlighted in yellow and β-strands in grey. TMHs and β-strands in *B. subtilis* KimA are numbered. Residues involved in potassium binding are indicated with a red asterisk. Residues implicated in c-di-AMP binding and inhibition (see text) are highlighted in cyan.

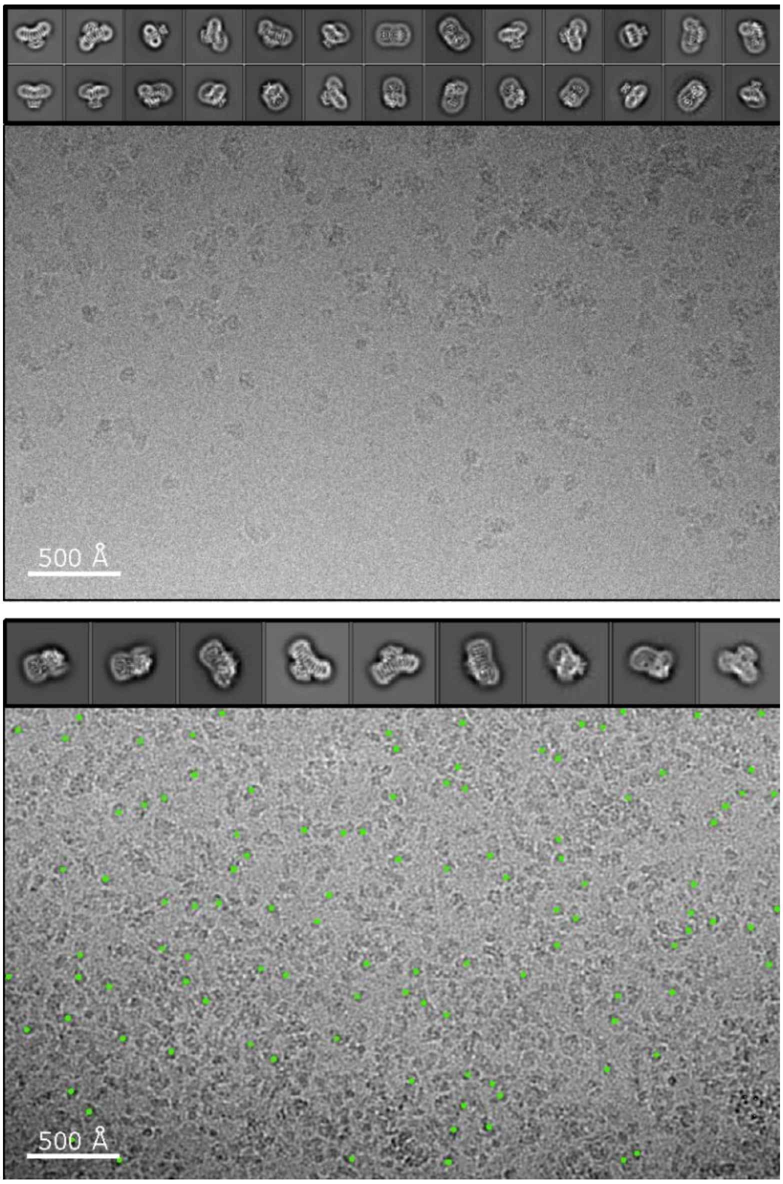


**Supplementary Figure 7:** Exemplary cryo-electron micrographs of KimA in DDM with a ten-fold excess of c-di-AMP (top) and reconstituted in amphipols (bottom, best picks marked green), respectively, and best 2D class averages.

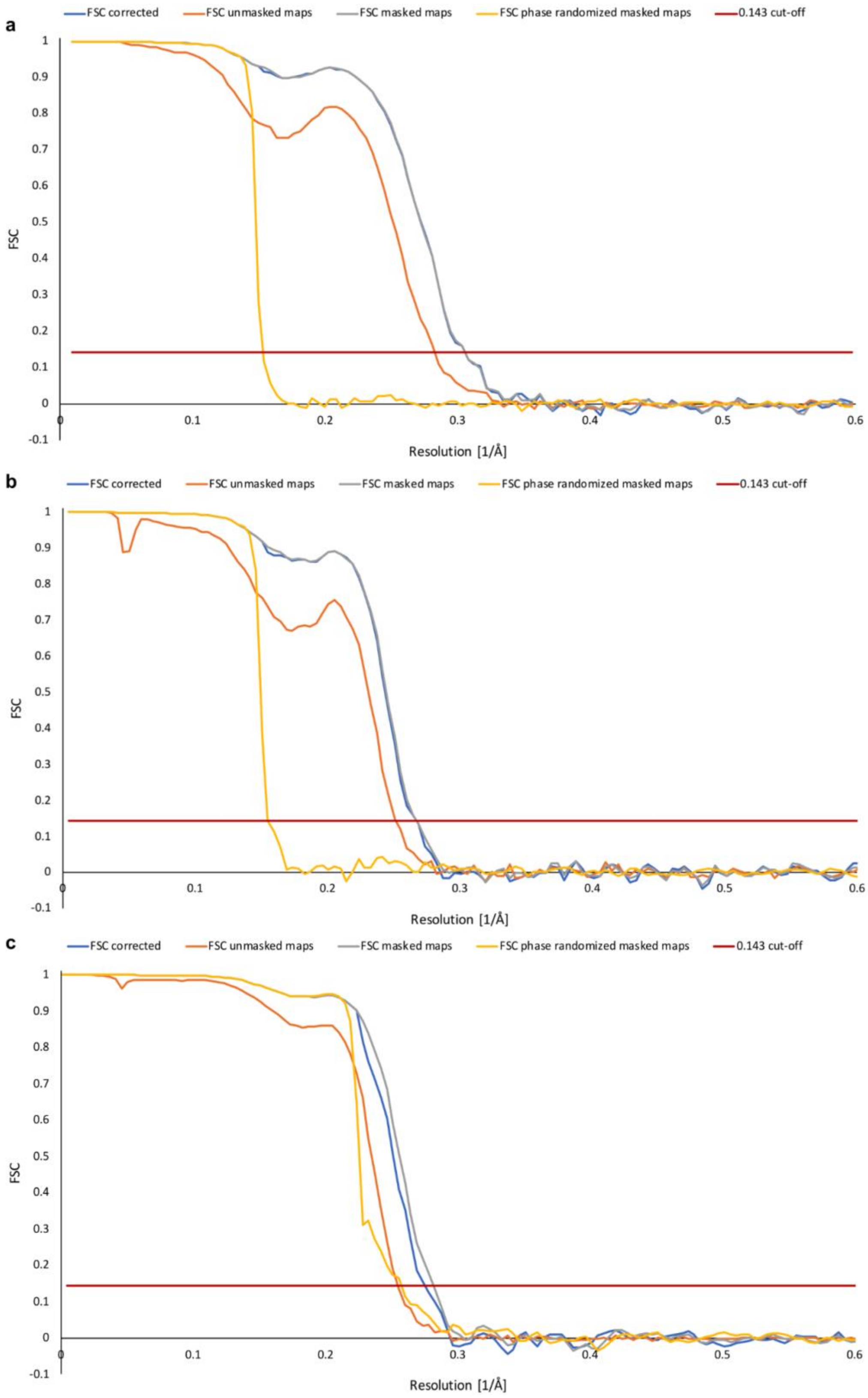


**Supplementary Figure 8:** FSCs of final maps of KimA with bound c-di-AMP, determined by the postprocessing procedure in Relion. **a)** C2 refinement of 332k particles of KimA solubilized in DDM with ten-fold excess of c-di-AMP added after purification. **b)** 296k particles of KimA in amphipols with C2 symmetry applied during refinement. **c)** 307k C2-symmetry expanded particles (KimA in amphipols) after focused 3D-classification and 3D-refinement focused on one half of the dimer.

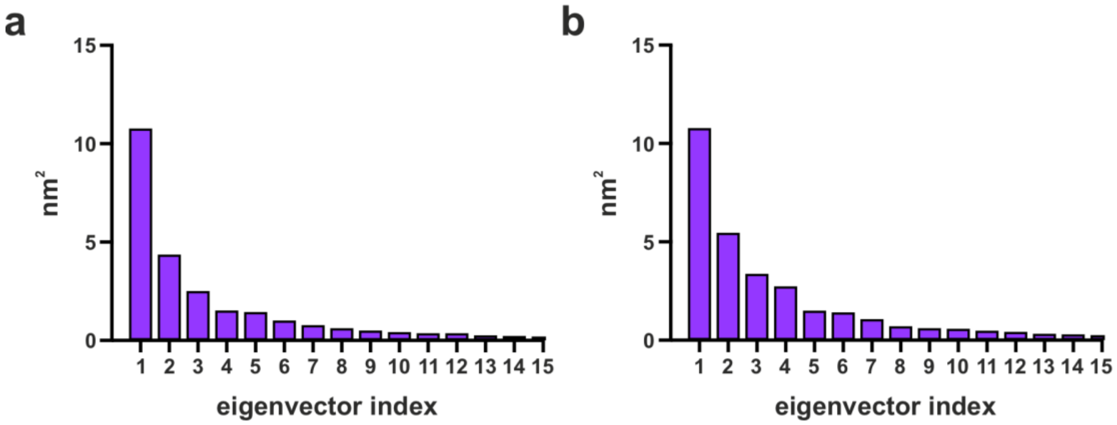


**Supplementary Figure 9: Contribution of each eigenvector to the total variance (nm$^2$).** Only the top 15 are shown out of ca. 1700. In both cases, eigenvector 1 and 2 account for ca. 50% of the total variance. **a)** Ligand, **b)** No ligand.

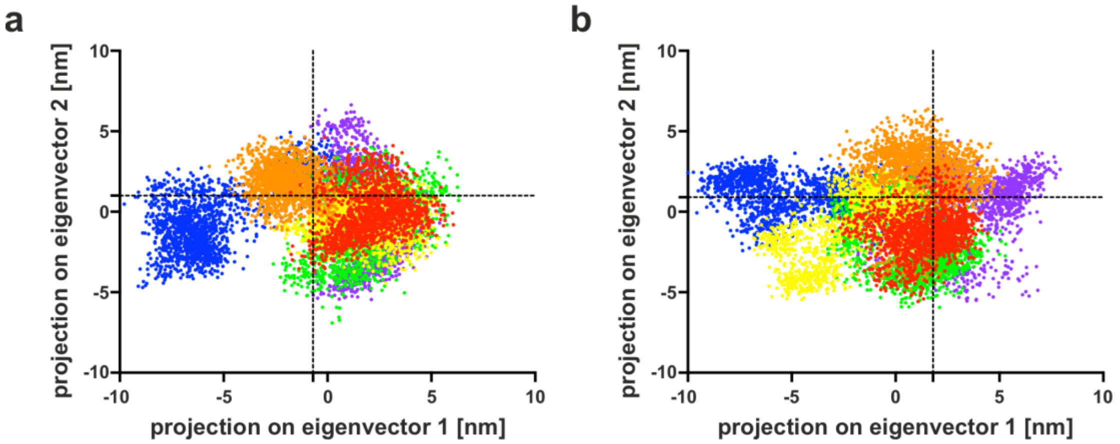


**Supplementary Figure 10: 2D projection of eigenvectors 1 and 2 for all 6 monomers (in 3 simulations).** The dotted lines represent the position of the structure in relation to the eigenvectors. **a)** Ligand **b)** No ligand.

**Supplementary Table 1: Primer used for site-directed mutagenesis in pB24*kimA*.**

| **Primer name** | **Sequence 3'□5'** |
|---|---|
| **KimA R337A Fwd** | CCAGTATATGCCGGCAATGTTTACAGTCAG |
| **KimA R337A Rev** | CCCTGACTGTAAACATTGCCGGCATATACT |
| **KimA A481W Fwd** | GTTGTGATTGTGCCTGTGTGGGGTGTCACCACCG |
| **KimA A481W Rev** | CGGTGGTGACACCCCACACAGGCACAATCACAAC |
| **KimA S582W Fwd** | CCTTCACAACCAATGGGCCTTCCTCCTCAGAG |
| **KimA S582W Rev** | CTCTGAGGAGGAAGGCCCATTGGTTGTGAAGG |
| **KimA Y118A Fwd** | GCTTGTTGATGCTATTTTAACAGTAG |
| **KimA Y118A Rev** | CTACTGTTAAAATAGCATCAACAAGC |
| **KimA N237A Fwd** | GAGGCCATTTCTGCTGCGATTCCTG |
| **KimA N237A Rev** | CAGGAATCGCAGCAGAAATGGCCTC |

**Supplementary Table 2: Cryo-EM data collection, refinement and model statistics**

| | DDM | Amphipols |
|---|---|---|
| **Data collection**<br>Microscope | FEI Titan Krios | FEI Titan Krios |
| Camera | Gatan K3 BioQuantum | Gatan K3 BioQuantum |
| Voltage (kV)<br>Nominal magnification<br>Calibrated pixel size (Å) | 300<br>105,000x<br>0.837 | 300<br>105,000x<br>0.837 |
| Electron exposure ($e^-$/Å$^2$) | 55.0 | 55.0 |
| Exposure time total (s)<br>Number of frames per image | 2<br>50 | 2<br>50 |
| Defocus range (µm) | -1.1 – -2.5 | -1.1 – -2.5 |
| **Image processing**<br>Motion correction software | *Relion* | *Relion* |
| CTF estimation software | *Gctf* | *Gctf* |
| Particle selection software | *Topaz* | *Topaz* |
| Micrographs (no.) | 2,349 | 4,303 |
| Initial particle images (no.) | 950,000 | 5,180,000 |
| Final particle images (no.) | 332,00 | 296,000 |
| Symmetry | C2 | C2 |
| Final resolution (Å) | 3.3 | 3.8 |
| **Refinement statistics**<br>Modeling software | *COOT, PHENIX* | *COOT, PHENIX* |
| Protein residues | 572 | 552 |
| Ligands | 2 c-di-AMP, 2 $K^+$,<br>8 DDM | 2 c-di-AMP |
| Map CC (volume) | 0.86 | 0.73 |
| RMS deviations<br>Bond lengths (Å) | 0.006 | 0.005 |
| Bond angles (°) | 0.740 | 0.613 |
| Average B-factor (Å$^2$) | 119.16 | 19.79 |
| Ramachandran plot<br>Outliers (%) | 0.0 | 0.0 |
| Allowed (%) | 5.0 | 5.0 |
| Favored (%) | 95.0 | 95.0 |
| Rotamer outliers (%) | 1.0 | 0.4 |
| Molprobity score | 1.77 | 1.95 |
| All-atom clashscore | 8 | 12 |

**Supplementary Videos**

V1: Cooperativity Network mobility of CDs in the absence of c-di-AMP. Movie showing the dynamics of selected residues in the KimA cytosolic domain in the absence of c-di-AMP. Movie made over a single 2.2 µs simulation trajectory using VMD.

V2: Cooperativity Network mobility of CDs in the presence of c-di-AMP. Movie showing the dynamics of selected residues in the KimA cytosolic domain in the presence of two bound c-di-AMP. Movie made over a single 2.2 µs simulation trajectory using VMD.

V3: Cooperativity Network mobility of CDs in the presence of only one. Movie showing the dynamics of selected residues in the KimA cytosolic domain in the presence of only one bound c-di-AMP. Movie made over a single 0.5 µs simulation trajectory using VMD.
